# Proteins and Ions Compete for Membrane Interaction: the case of Lactadherin

**DOI:** 10.1101/2020.04.03.023838

**Authors:** A. M. De Lio, D. Paul, R. Jain, J. H. Morrissey, T. V. Pogorelov

## Abstract

Charged molecular species, such as ions, play a vital role in the life of the cell. In particular, divalent calcium ions (Ca^2+^) are critical for activating cellular membranes. Interactions between Ca^2+^ and anionic phosphatidylserine (PS) lipids result in structural changes of the plasma membrane and are vital for many signaling pathways, such as the tightly regulated blood coagulation cascade. Upon cell damage, PS lipids are externalized to the outer leaflet, where they are not only exposed to Ca^2+^, but also to proteins. Lactadherin is a glycoprotein, important for cell-adhesion, that can act as an anticoagulant. While a number of experimental studies have been performed on lactadherin’s C2 domain’s (LactC2) binding affinity for PS molecules, an atomistic description of LactC2 interactions with PS lipids in the plasma membrane is lacking. We performed extensive all-atom molecular dynamics simulations of mixed lipid bilayers and experimental characterization of LactC2-membrane interactions in the presence and absence of Ca^2+^ and characterized PS-Ca^2+^ and PS-LactC2 interactions to guide our understanding of how these interactions initiate and impede blood coagulation, respectively. The captured spontaneously formed long-lived PS-Ca^2+^ and PS-LactC2 complexes revealed that the protein side chains involved in PS-LactC2 interactions appear to be affected by the presence of Ca^2+^. The degree of LactC2 insertion into the lipid bilayer also appears to be dependent on the presence of Ca^2+^. Characterizing the interactions between Ca^2+^ and LactC2 with PS lipids can lead to a greater understanding of the activation and regulation of the blood coagulation cascade and of the basis of charged species interactions with the lipid membrane.

**STATEMENT OF SIGNIFICANCE:** Lactadherin plays an important role in cellular signaling including blood coagulation. Many of these processes involve lactadherin interacting with the lipids of the cell plasma membrane. Lactadherin acts as an anticoagulant and contributes to a number of health issues. Understanding the interactions that drive lactadherin’s anticoagulant properties can lead to potential new drug targets.

## INTRODUCTION

Lactadherin, originally identified as a milkfat globule glycoprotein, has been shown to bind to the plasma membrane via interactions with phosphatidylserine (PS) lipids. Lactadherin has been found to participate in a number of cellular processes including bilayer stabilization (1), mediating apoptotic clearance of lymphocytes (2, 3) and epithelial cells (4), and clearing amyloid ß-peptide, a key component of Alzheimer disease. Lactadherin has also been found to help regulate the blood coagulation cascade by acting as an anticoagulant (5).

Bovine lactadherin, studied here, is comprised of four domains, two epidermal growth factor domains and two discoidin domains (EGF1-EGF2-C1-C2). The C2 domain of lactadherin (LactC2) (Fig. 1) has been found to be the primary location of PS binding (6).

**Figure 1.**
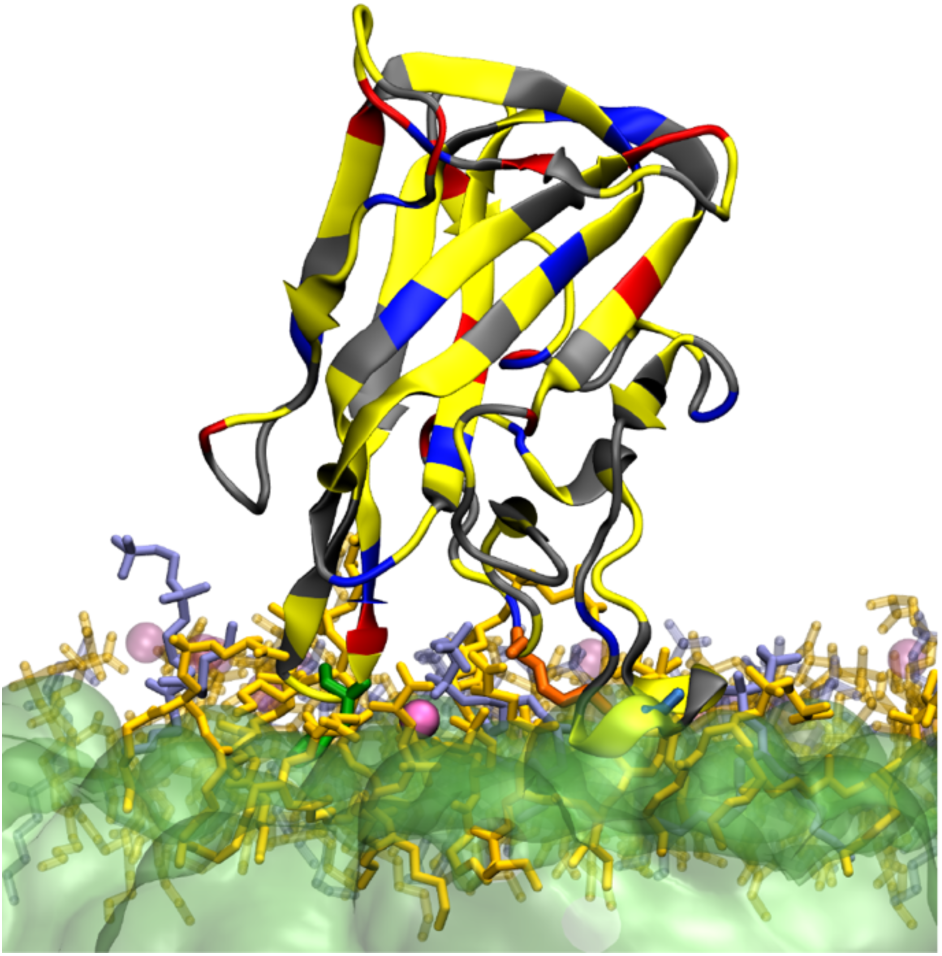
Snapshot of LactC2 inserted in the short-tail PS (orange)/PC (gray-blue) HMMM plasma membrane with dichloroethane (DCLE) (green) solvent in the presence of Ca^2+^ (pink). Transparent lipids and Ca^2+^ are farther than 3Å from LactC2 residues. Residues colored by type: polar (gray), non-polar (yellow), basic (blue), and acidic (red).

LactC2 functions as an anticoagulant by blocking PS binding sites from use by coagulation factors such as factor VIII, the prothrombinase complex, and the factor VIIa-tissue factor complex, many of which are mediated by divalent calcium ion (Ca^2+^)-PS interactions. Unlike many of the components of the coagulation cascade, LactC2 does not require Ca^2+^ to mediate interactions with the membrane surface (5). This suggests that a direct competition could exist between LactC2 and Ca^2+^ for PS binding sites.

The LactC2 domain is structurally similar to the C2 domains in some blood coagulation factors, such as factor V (fV-C2) and factor VIII (fVIII-C2). LactC2, fV-C2, and fVIII-C2 all contain long loops, here referred to as Spikes, that are comprised of water-exposed hydrophobic residues. Site-directed mutagenesis of these hydrophobic residues has highlighted their importance in the functionality of lactadherin. It has been hypothesized that the hydrophobic residues of these Spikes insert into the membrane (7), but detailed studies are lacking. Characterizing the types of interactions that take place at the membrane surface can aid in our understanding of lactadherin’s role as a coagulation cascade regulator and, more generally, how charged molecular moieties affect and shape the plasma membrane.

Not only do the interactions between the spikes of LactC2 and the membrane affect the coagulation cascade, they also dictate the formation of aortic medial amyloids (AMA), which are present in 97% of the Caucasian population over 50 years of age (8) and may be linked to thoracic aneurysms (9, 10). Since the peptide forming the AMAs, medin, is derived from the region spanning Spikes 2 & 3 of the human LactC2 domain, characterizing the dynamics and interaction potentials between lactadherin and the plasma membrane are critical for elucidating the biochemical properties of lactadherin.

Here we utilize all-atom molecular dynamics (MD) simulations and experimental binding studies to capture LactC2 insertion into and interactions with the plasma membrane. Characterization of key charged species-lipid interactions both in the presence and absence of Ca^2+^ are also presented.

## MATERIALS & METHODS

### Molecular Dynamics Simulations

#### System Preparation

The membrane system studied consisted of a lipid bilayer of 120 lipids with a 70:30 palmitoyloleoylphosphatidylserine: palmitoyloleoylphosphatidylcholine, POPS:POPC, molar ratio and an approximate initial density of 60 Å^2^ per lipid. The choice of fluid POPS and POPC in favor of more rigid phospholipids was made to the increase membrane association and insertion (11). The use of a 70% PS lipid bilayer, while not physiologically accurate, was used to ensure capture and extensive sampling of LactC2-PS interactions. The independently prepared replicas of the lipid bilayer were initially modeled with the Highly Mobile Membrane Mimetic (HMMM) model (12–14) which consisted of the hydrophobic organic solvent 1,1-dichloroethane (DCLE) between two leaflets of the short-tail versions of the POPS and POPC lipids in the bilayer. The use of HMMM membranes increases lipid lateral diffusion in the membrane by 1-2 orders of magnitude, while preserving all-atom description. The increased lateral diffusion subsequently increases the sampling rate of lipid-lipid and lipid-ion interactions (12, 13, 15, 16). The HMMM membranes were assembled in the CHARMM-GUI (17–19) web portal with the HMMM Builder module (20). Divalent calcium ions were added to the system on both sides of the lipid bilayer in a concentration of 5mM to mimic the lipid:ion ratio of the nanodiscs used in SPR experiments. In each replicate, a single copy of the bovine LactC2 domain (PDB ID: 3BN6) (7) was added approximately 20 Å above the membrane surface. The resultant membrane:ion:protein systems were solvated with ∼27,000 water molecules. Ten independent replicates containing Ca^2+^ and ten control replicates with no Ca^2+^ were prepared. After the HMMM MD simulations, the HMMM Builder module (20) in the CHARMM-GUI (17–19) web portal was used to convert the short-tail lipids into full-tail lipids and remove the DCLE solvent between the leaflets.

#### MD Simulation Protocols

MD simulations of both systems were performed using NAMD 2.12 (21). CHARMM36 force field (22) parameters were used for the LactC2, lipids, and ions, while the CGenFF (23) force field was used for DCLE molecules, and water molecules were modeled by the TIP3P (24) model. The HMMM systems were simulated for 250 ns. After lipid tail conversion, the systems were simulated for an additional 100 ns, and the latter were used for analyses. All simulations were performed at a constant temperature of 303 K and at a constant 1 atm of pressure. Constant pressure was maintained by the Langevin piston Nose-Hoover method (25, 26) with a damping coefficient of 0.5 ps^-1^. The nonbonded cutoff distance for short-range interactions was 12 Å with switching at 10 Å. The particle mesh Ewald (PME) method (27, 28), with a 1 Å grid spacing, was used for long-range electrostatics. Hydrogen atom bond lengths were restrained with SETTLE (29). The integration step was set to 2 fs.

#### Analysis of MD Simulations

Statistical analysis of the MD trajectories was performed using version 3.7 of the Python programming language (30). The matplotlib (31), seaborn (32), pandas (33), NumPy (34), and MDAnalysis (35, 36) libraries were utilized in the analysis of the protein insertion and residue-lipid contacts via the Spyder (37) interface. Statistical analysis of both systems was performed on all replicates were LactC2 bound to the membrane. In every simulation the LactC2 domain spontaneously interacted with the lipid bilayer, although in one replicate of the Ca^2+^ containing system, only Spike 1 interacted with the membrane as the domain interacted laterally with the membrane. To monitor convergence of systems, three residues, one each from Spikes 1-3, were monitored during the combined HMMM and full-tail trajectories for their depths of insertion into the membrane (Figs. 2, S2, S3).

**Figure 2.**
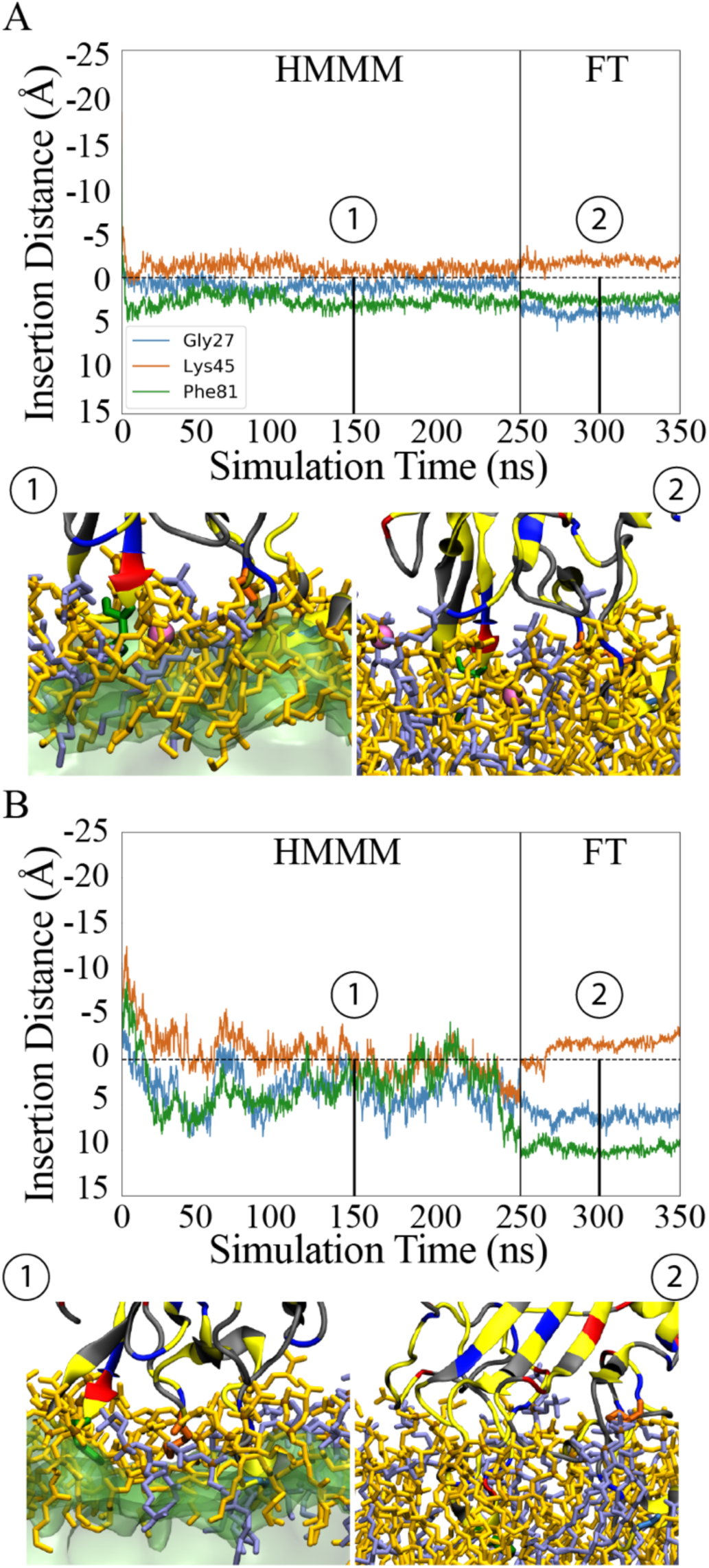
Convergence of LactC2 Spike residues insertions into HMMM (0-250 ns) and full-tail (250-350 ns) lipid membranes for one replicate of each system: with Ca^2+^ (A) and without Ca^2+^ (B). Snapshots are taken from 150 ns (HMMM) and 300 ns (full-Tail).

Insertion of protein residues into the lipid membrane was determined by creating a plane at the level of the phosphorous atoms of both the POPS and POPC lipids. A residue was defined to be inserted into the membrane when its center of mass moved past this plane, deeper into the membrane. The interactions between protein residues, both in and near the surface of the membrane, and the charged lipid headgroups were analyzed using contact number with the cutoff distance of 5 Å. The mean contact number represents the number of lipid phosphate groups contacting each residue at all points during the analysis trajectory, averaged over the number of replicates.

The NetworkView (38, 39) plugin of VMD (40) was used for the dynamical network analysis of LactC2. One replicate from both systems was used for network analysis. In addition, one copy of LactC2 was placed in a water box and simulated for 100 ns. For the membrane systems, the full 100 ns of full-tail simulation data were used, and for the water box system, the whole 100 ns simulation was used for analysis. All network analysis was performed on the trajectories with a time step of 100 ps. The NetworkView (38, 39) plugin was also used to perform the community analysis via the Girvan-Newman algorithm (41).

## Experimental Methods

### Materials

POPC and POPS lipids, purchased from Avanti Polar Lipids (Alabaster, AL), were used to prepare the nanodiscs used in this study. Bio-Beads® SM-2 adsorbent was purchased from Bio-Rad. Bovine Lactadherin was purchased from Hematologic Technologies (Essex Junction, VT). For the Biacore binding studies we purchased the series S Nitrilotriacetic acid Biacore sensor chips from GE Healthcare. The recombinant His-Tagged MSP was expressed in *Escherichia coli* BL21 DE3 cells (Agilent technologies) and purified in AKTA start system (GE Healthcare Life Science) by Ni NTA(Qiagen) column.

### Preparation of Nanodiscs

Quantification of the binding constants for lactadherin to PS containing membranes were done with nanodiscs. Nanodiscs are soluble monodisperse discoidal lipid/protein particles with controlled size and phospholipid composition where the lipid bilayer is surrounded by a helical protein belt termed as membrane scaffold proteins (MSP) (42). To prepare the nanodiscs, a total of 3.9 μmol of phospholipids comprised of 70% POPS and 30% POPC were prepared in chloroform and dried under compressed nitrogen. Nanodiscs comprised of 100 % PC were also prepared as a negative control for the binding studies. The residual chloroform from the lipids was removed overnight under high vacuum. The lipids were dissolved in a 100 mM sodium deoxycholate, 20mM Tris, and 100mM NaCl buffer solution and incubated with MSP for 2 hrs at 4°C. The detergent was removed post incubation via Bio-Beads at 4°C for 4 hours in a rotator. This allowed the self-assembly of the nanodiscs. The nanodiscs were further purified by size exclusion chromatography (SEC) in pure AKTA with the Superdex 200 10/300 GL column (GE healthcare Life Sciences).

### Surface Plasmon Resonance Analysis of Lactadherin binding to Nanoscale Bilayers

The binding affinities for lactadherin to the nanodiscs in the presence and absence of 5 mM Ca^2+^ were quantified by surface plasmon resonance (SPR) on a GE Biacore T-200 instrument. His-Tagged nanodiscs were loaded on the Ni-NTA surface sensor chip in a solution of 20 mM HEPES and 100 Mm NaCl. The flow rate of lactadherin was set to 30 uL/min and the binding was monitored in real time as the concentration of lactadherin was increased from 25 nM to 2.5 µM. The surface equilibration time for the binding studies was 110 seconds. The protein concentration used to study membrane association times both with and without Ca^2+^ was 800 nM. Experiments were completed in triplicate with fresh samples.

As reported previously (43), the binding isotherms were plotted from maximal steady-state response units versus the protein concentration flowed over the chip surface, from which the equilibrium binding constant, K_d_, values were derived by fitting the single-site ligand binding equation to the data in Graph Pad Prism 8.0 (44). The single-site ligand binding equation is:

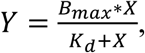

where B_max_ is the maximum specific binding,

X is the concentration of lactadherin,

K_d_ is the equilibrium binding constant.

Specific binding of lactadherin was corrected by normalizing it to the 100% PC nanodiscs. Analysis of the association rate, k_on_, values were performed in accordance with the protocol established by Qi, et al. (45) (Fig. S8) using the equation:

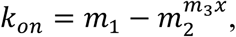

where m_1_ is the initial value of response in the association phase,

m_2_ is the final value of response in the association phase,

m_3_ is the exponential value of the association phase.

The binding assay setup used here has the mass transport effect limitation, which is nullified during the steady state analysis.

## RESULTS & DISCUSSION

### Membrane Insertion

To investigate the prevalent types of interactions between LactC2 and the plasma membrane, it was necessary to capture the protein-membrane interface at the atomic resolution. Crystal structure analysis of the LactC2 domain suggests that Spikes 1 and 3 can insert into the plasma membrane as they contain a number of hydrophobic residues (7). To capture and characterize insertion, two systems were assembled, one containing a physiologically relevant 5 mM concentration of Ca^2+^ and one in the absence of Ca^2+^ (Fig. S1). We performed extensive, total of 7 μs, all-atom equilibrium MD simulations using the HMMM (13, 14, 46) and full-tail lipid models of both systems and observed reproducible spontaneous insertions of LactC2 into the lipid bilayer (Figs. 1 & 3).

We observed consistent interactions of Spike 1, 2, & 3 with the bilayer in the presence (Figs. 3A and S4), and absence (Figs. 3B and S5) of Ca^2+^. The height of the bars in Fig. 3 indicate the three quartiles of up- and-down motion experienced by the Spikes during the analyzed eplicate trajectories. The insertion data offers a number of insightful details about how the insertion of LactC2 is affected by the presence of Ca^2+^. The degree of vertical motion experienced by LactC2 is statistically similar independent of the presence of Ca^2+^ (Figs. 3, S4 and S5). Since lactadherin is able to bind to the plasma membrane without a mediator (5), it was anticipated that once spontaneous interactions occurred between LactC2 and lipid headgroups, that the domain would not move far away. We also observe that the depth of insertion of the Spike 2 can be greater in the presence of Ca^2+^ (Figs. 3, S4, S5). The charged residues K24 and K45 stop just at, or very near, the surface of the membrane in the absence of Ca^2+^ but can insert up to 6 Å in the presence of Ca^2+^ (Figs. S4D&E). The ability of the charged Spike residues to insert into the bilayer, rather than just anchor on top, subsequently allows the hydrophobic residues of the Spikes to insert more deeply into the hydrophobic core of the bilayer and anchor the domain. One hypothesis is that the Ca^2+^ moieties cluster the PS lipids (47, 48), thus removing some possible instances of the positively charged lysine residues interacting with the negatively charged PS headgroups. The opposite phenomenon is also observed for residue D80 (Fig. 3). In the systems containing Ca^2+^, the negatively charged D80 has the potential to interact with the positively charged Ca^2+^ moieties, which inhibit Spike 3 from further inserting into the membrane. These D80-Ca^2+^ interactions and subsequent stunted insertion is observed is several Ca^2+^ containing replicates (Figs. 4C & S4).

**Figure 3.**
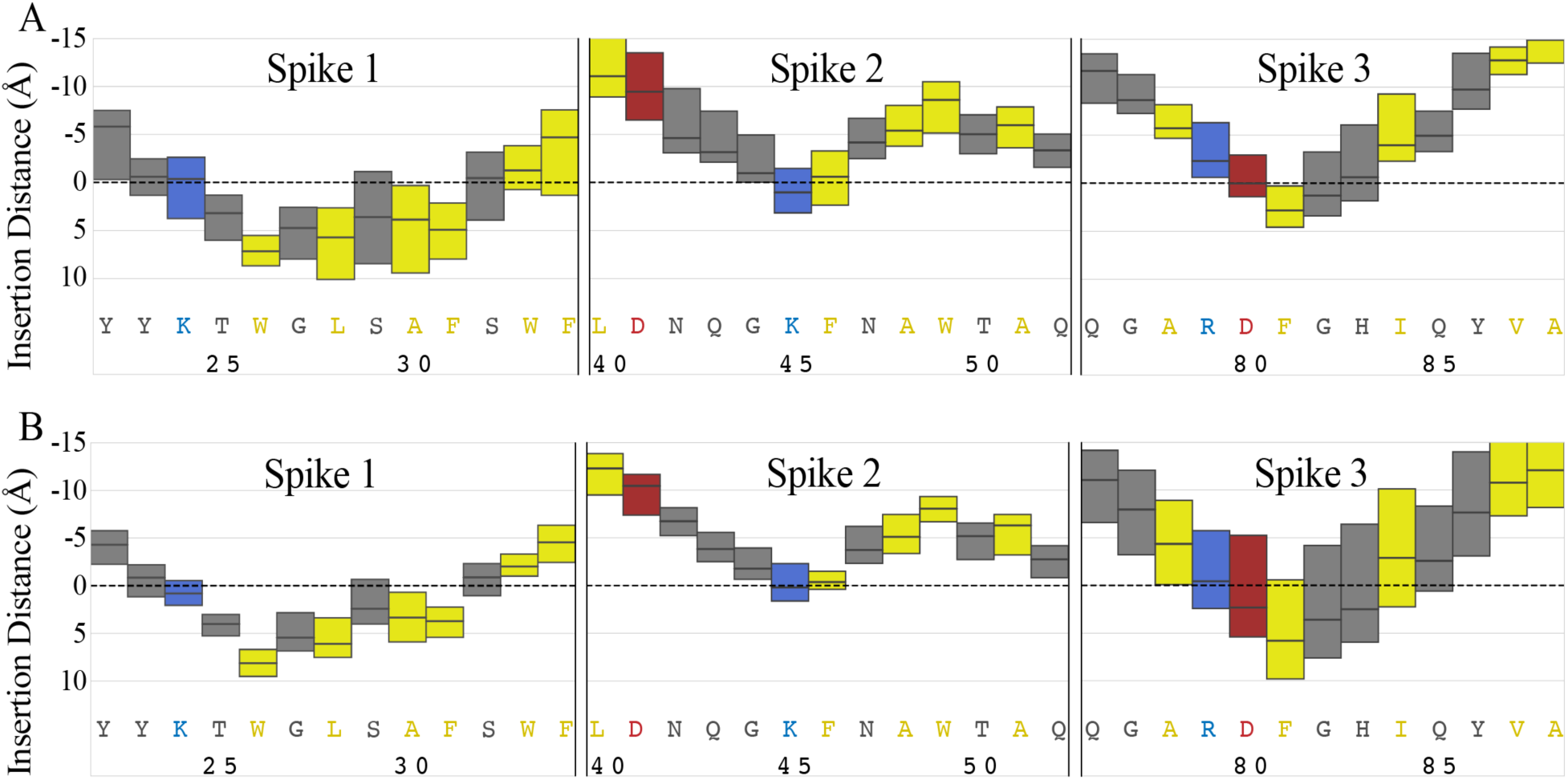
Average insertion of Spikes 1, 2, 3 of LactC2 into membrane. Insertion distance between the lipid phosphorous atom plane (dashed line) and the center of mass of the residues over 100 ns of full-tail lipid MD simulations. Residue insertion depths are positive inside the membrane and are color coded by residue type: polar (gray), non-polar (yellow), basic (blue), and acidic (red). (A) Insertion of LactC2 residues in the replicas containing Ca^2+^. (B) Insertion of LactC2 residues in the replicas containing no ions. Three quartiles are shown as box plot.

**Figure 4.**
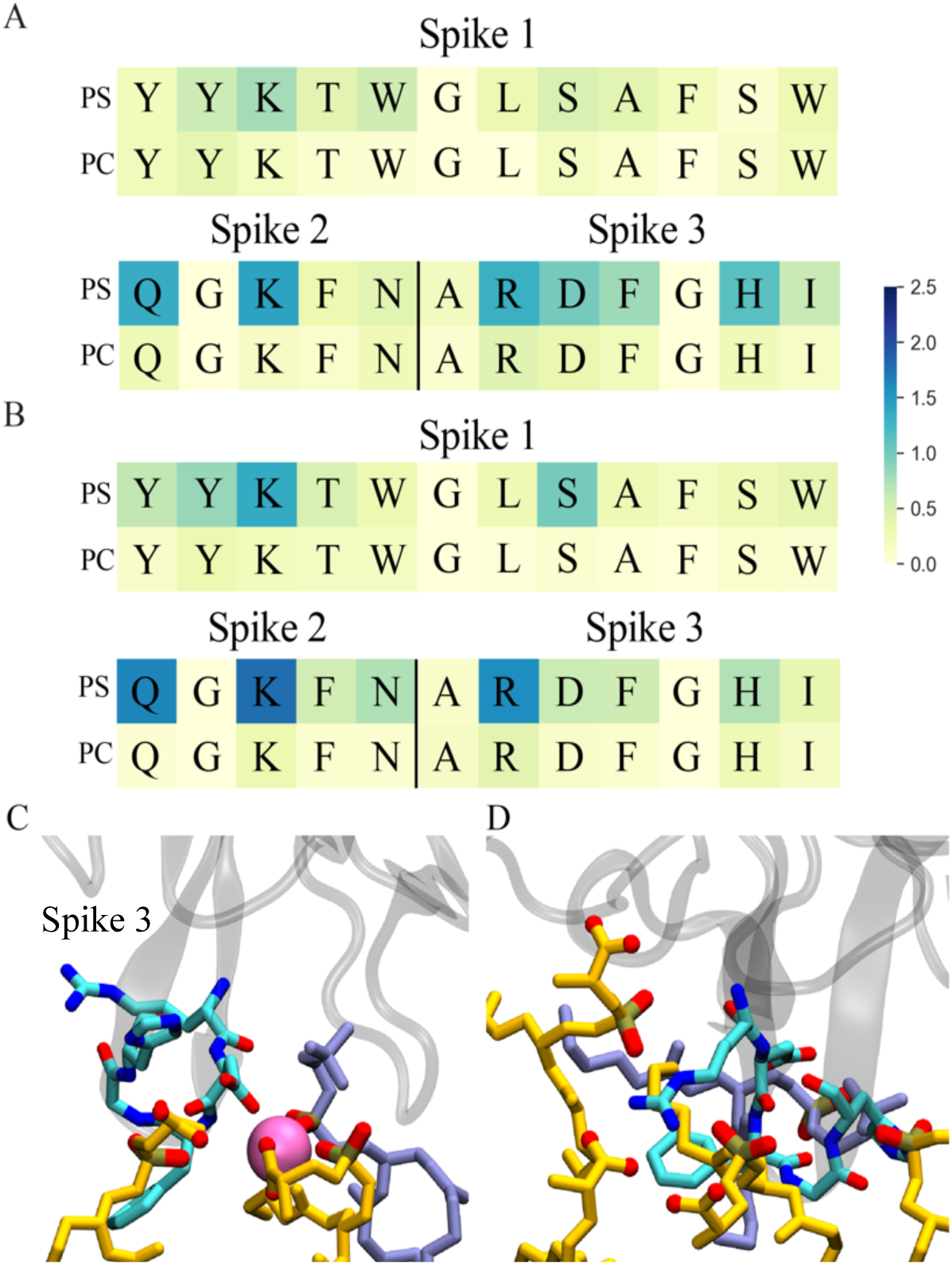
Mean contact number between the residues in Spikes 1-3 of LactC2 and individual PS and PC headgroups in the membrane (A) in the presence of Ca^2+^ and (B) in the absence of Ca^2+^. Snapshots of Spike 3 residue interactions (C) in the presence of Ca^2+^ and (D) in the absence of Ca^2+^. Residues are colored by atom name: carbon (cyan), nitrogen (blue), and oxygen (red). Lipids PS (orange) and PC (gray-blue) show headgroup carboxy oxygens (red) and phosphate groups colored by atom type.

We took a closer look at the number of PS and PC contacts the Spike residues encountered both in the presence and absence of Ca^2+^ (Figs. 4A and B). We also observe that, in both systems, LactC2 has a larger number of contacts with PS than with PC at any one time. This is consistent with previous studies which suggest that LactC2 preferentially binds PS and does not bind PC (6, 49). A lack of interaction with PC headgroups is not unknown, especially in the blood coagulation cascade. Previously reported, our “Anything But Choline” (ABC) hypothesis suggests that the bulkiness of the PC headgroup inhibits interactions with GLA domains and enzymes (50). In the absence of Ca^2+^, positively charged residues, such as K45 in Spike 2, have more simultaneous interactions with multiple PS lipids (Fig. 4B). Fig. 4D shows a snapshot of the interactions between R79 and nearby PS lipids. In similarity to the interactions between Ca^2+^ and PS lipids (47), we find both the carboxy oxygens and phosphate oxygens of PS interact with R79.

Several instances of LactC2-Ca^2+^ interactions were observed, which demonstrates that LactC2 residues can also interact with Ca^2+^ moieties via sidechain carboxy oxygens (Fig. 4C). This direct LactC2-Ca^2+^ interaction was not anticipated as LactC2 does not functionally require Ca^2+^ (5), however based on the electrostatic properties of D80 and Ca^2+^, it is not surprising.

To determine the effect interactions have on the stability of the LactC2 secondary structure, we performed network analysis on one LactC2 replicate in both systems and on a LactC2 copy in a water box (Fig. S6). The edges (lines) between nodes (spheres) in Spike 3 show the greatest variance in the different environments. In the water box system (Fig. S6C), the edges that join the nodes in Spike 3 are thickest, indicating strong interactions between the residues in that region. When Spike 3 interacts with the lipid bilayer (Fig. S6A&B), the thickness of the edges decreases, Indicating that the interactions destabilize the structure of Spike 3.

Surface plasmon resonance (SPR) studies of lactadherin on PS/PC nanodiscs were performed in order to determine whether the presence of Ca^2+^ moieties affects the binding affinity (Figs. 5A & 5C) and association time (Fig. 5B & S6) of lactadherin to the membrane surface. To our knowledge, this is the first report on the binding affinity of lactadherin using nanodiscs as the membrane model. We find that the binding affinity in the absence of Ca^2+^ is 296.2 ± 82.8 nM, which is in agreement with previously reported literature (43). In order to more accurately model a physiologically relevant system, we also measured the binding affinity of lactadherin in the presence of Ca^2+^; which was measured to be 283.0 ± 66.5 (nM). These values suggest that there is no change in the binding affinity of lactadherin to the membrane model in the presence and absence of Ca^2+^ (Fig. 5A). Since LactC2 binds PS in a Ca^2+^-independent manner (5), we expected to observe similar binding affinities in both systems. This is also in agreement with our computational observation that LactC2-PS interactions occur regardless of the presence or absence of Ca^2+^ (Figs. 2 & 3).

**Figure 5.**
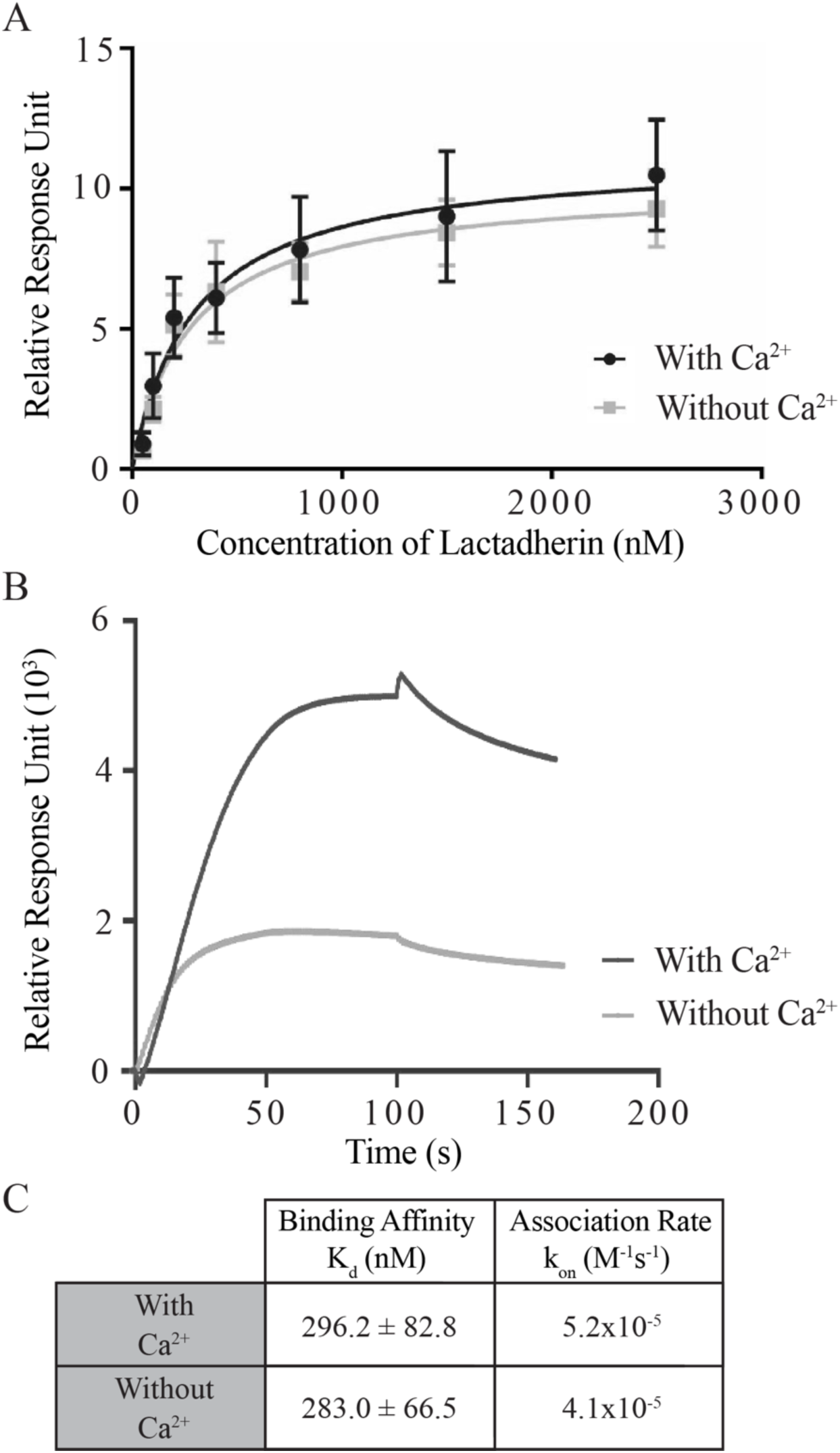
Experimental studies on lactadherin binding to PC/PS nanodiscs. (A) Comparison of binding affinities of lactadherin to nanodiscs containing 70% PS and 30% PC in the presence of (black) and absence of (gray) Ca^2+^ at multiple lactadherin concentrations. (B) Association times of lactadherin in the presence of (black) and absence of (gray) Ca^2+^ with a lactadherin concentration of 800 nM. (C) Binding affinities and association rates for both systems. The k_on_ was calculated using all the concentrations (Fig. S8).

We hypothesized that Ca^2+^ and the charged residues of the Spikes would compete for the charged lipid headgroup binding sites of the membrane. While we did not observe any exchange of PS binding sites between Ca^2+^ and LactC2 residues, likely due to the proportionally large concentration of PS in the model lipid bilayer, we used SPR to determine if the association time and association rates of lactadherin to the membrane were affected by the presence of Ca^2+^ (Fig. 5B). The k_on_ values for lactadherin in the presence and absence of Ca^2+^ were not significantly different enough to suggest competition between lactadherin and Ca^2+^ (Figs. 5C & S7), and therefore suggest that lactadherin’s anticoagulant properties do not simply stem from this type of competition. Instead, lactadherin’s anticoagulant properties could stem from its binding affinity to the membrane. In a 50% PS and 50% PC nanodisc, the K_d_ of factor X was reported as 380 ± 5 nM (51). Since the binding affinities for factor X and lactadherin (Fig. 5C) are similar, competition assays in consistently composed nanodiscs can be used to determine the degree of competition between them, but that is beyond the scope of this paper.

## CONCLUSIONS

We have captured the spontaneous interaction with and insertion into the lipid bilayer of the LactC2 domain. While the hydrophobic residues of the Spikes are integral for lactadherin function, the charge-driven interactions between the lipid headgroups and LactC2 or Ca^2+^ determine insertion depth, the mobility of LactC2 once bound to the membrane, and the stability of the backbone interactions. Novel binding studies of lactadherin to nanodiscs demonstrated that its binding affinity and association time to the membrane are independent of the presence of Ca^2+^. While it is known that lactadherin acts as an anticoagulant, our results suggest that its anticoagulant properties are not simply a result of PS site competition with Ca^2+^, but could be related to competition with blood clotting factors for membrane binding sites, based on membrane binding affinity data (51). This study provides new insight into the mechanism that drives the anticoagulant properties of lactadherin and will guide our future studies into determining lactadherin’s mechanism of anticoagulation, characterizing the effect the charged residues on Spikes 1-3 have on the shape and functionalization of the lipids in the membrane, and determining the functionality of the LactC2-derived medin amyloid fibrils.

## AUTHOR CONTRIBUTIONS

A.M.D. performed MD simulations on all 10 replicates of the system containing no Ca^2+^ moieties. R.J. performed MD simulations on the first 5 replicates of the Ca^2+^ containing system, and A.M.D. performed the MD simulations of the last 5 replicates of this system. A.M.D. performed all analysis of MD simulations including insertions depth, contact number, and network analysis. T.V.P. supervised the entirety of the computational studies. D.P. performed all binding studies and completed all the analysis corresponding to those studies. J.H.M. supervised the binding studies.

## ACKNOWLEDGEMENTS

The authors are grateful for support from a National Institutes of Health Transformative Research Award (R01 GM123455) and NHLBI grant R35 HL135823. T.V.P. acknowledges support from the Department of Chemistry, the School of Chemical Sciences, and the Office of the Vice-Chancellor for Research (RSOCR Award 4703) at University of Illinois at Urbana-Champaign and computational resources from the Texas Advanced Computing Center (TACC) at the University of Texas at Austin made available from the Extreme Science and Engineering Discovery Environment (XSEDE) Grant TG-MCB130112, which is supported by National Science Foundation Grant ACI-1053575. A.M.D. is supported by a postdoctoral fellowship from the National Center for Supercomputing Resource and Cyprus Institute.

## SUPPORTING MATTERIAL RESULTS

**Figure S1.**
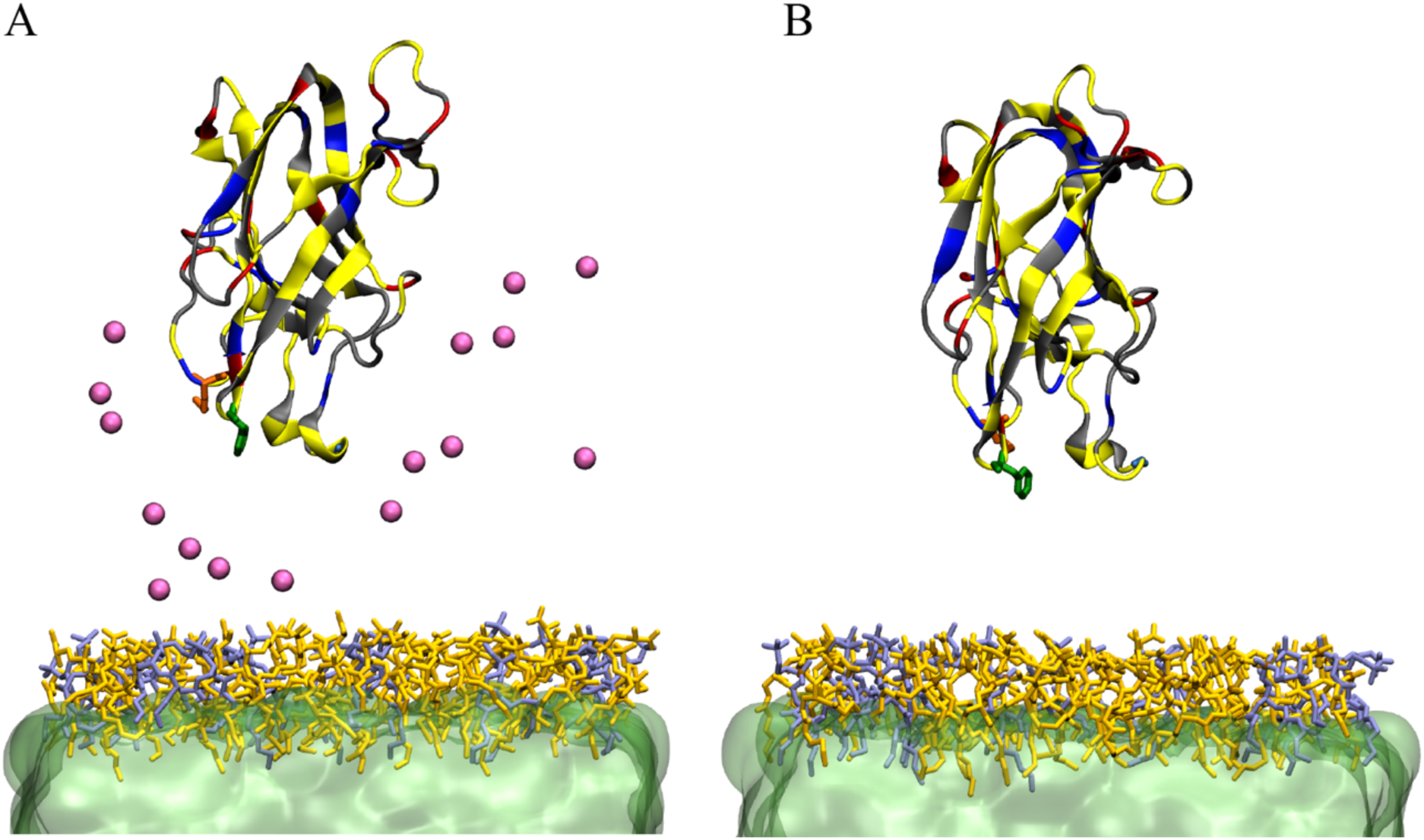
Snapshots of initial LactC2 systems (A) in the presence of Ca^2+^ and (B) in the absence of Ca^2+^ (pink spheres). Short-tail POPS (orange)/POPC (gray-blue) HMMM, plasma membrane with DCLE (green) solvent. Residues colored by type: polar (gray), non-polar (yellow), basic (blue), and acidic (red).

**Figure S2.**
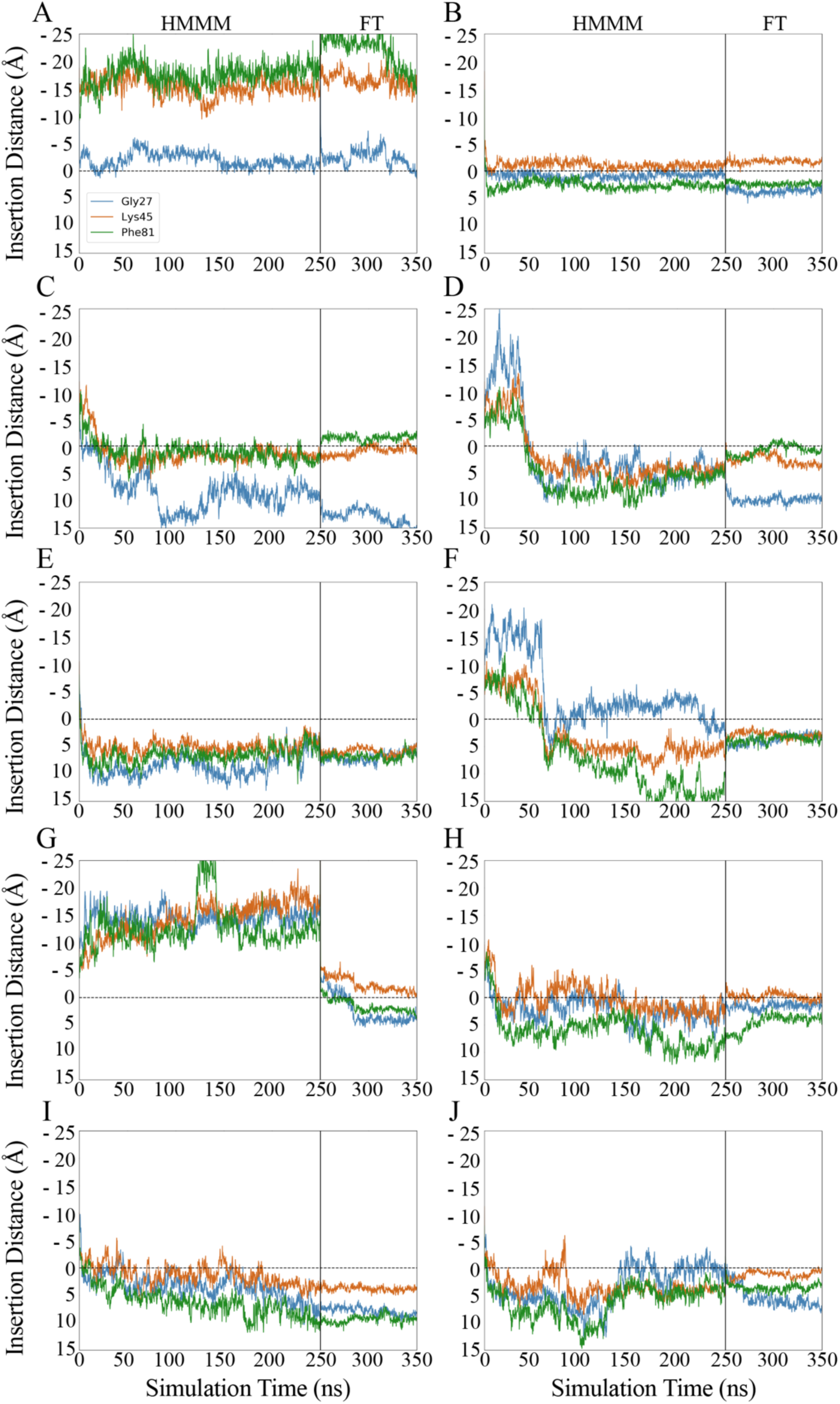
Partitioning of residues on Spikes 1-3 in HMMM and full-tail membranes for all 10 replicate systems in the presence of Ca^2+^ (A-J).

**Figure S3.**
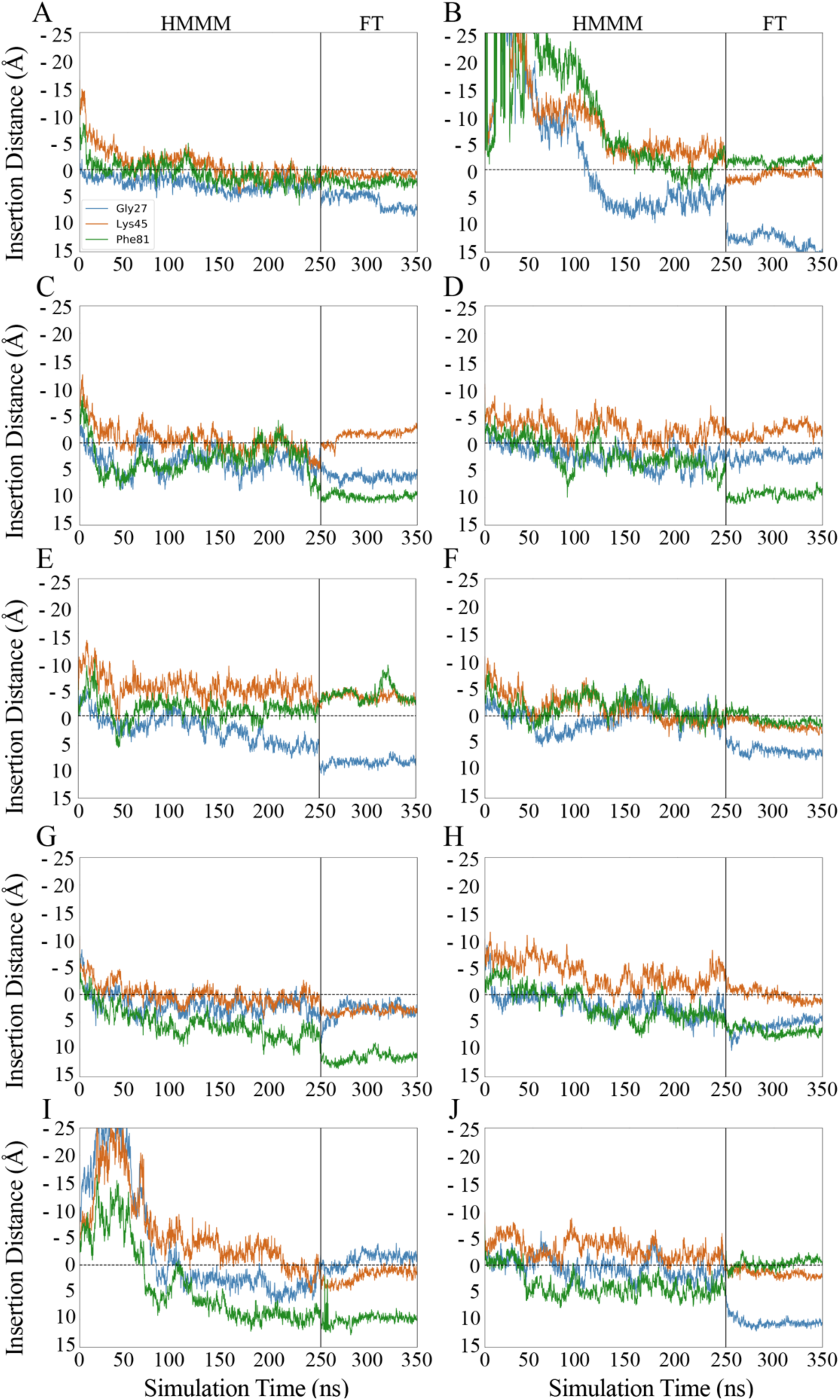
Partitioning of residues on Spikes 1-3 in HMMM and full-tail membranes for all 10 replicate systems without calcium ions (A-J).

**Figure S4.**
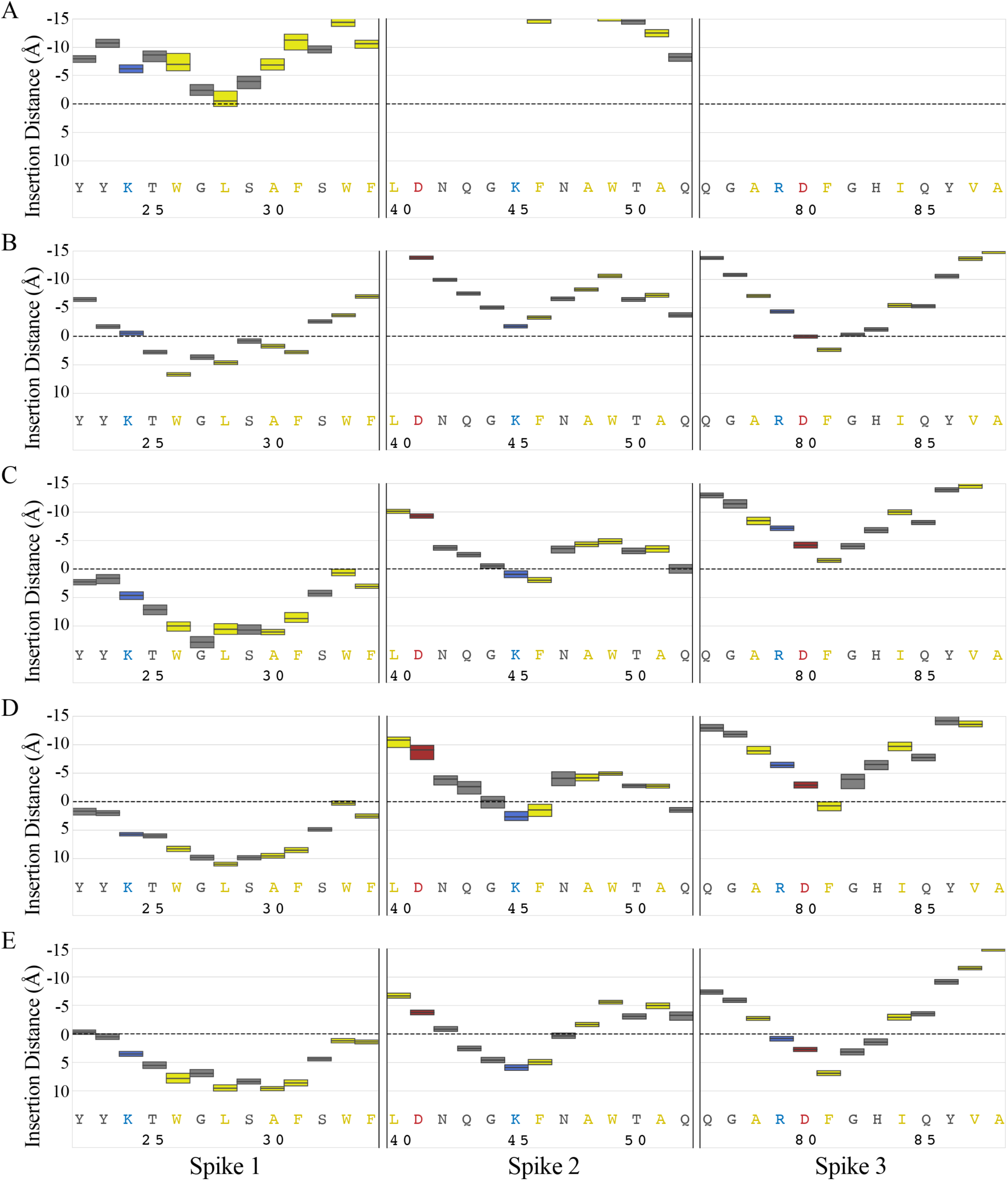
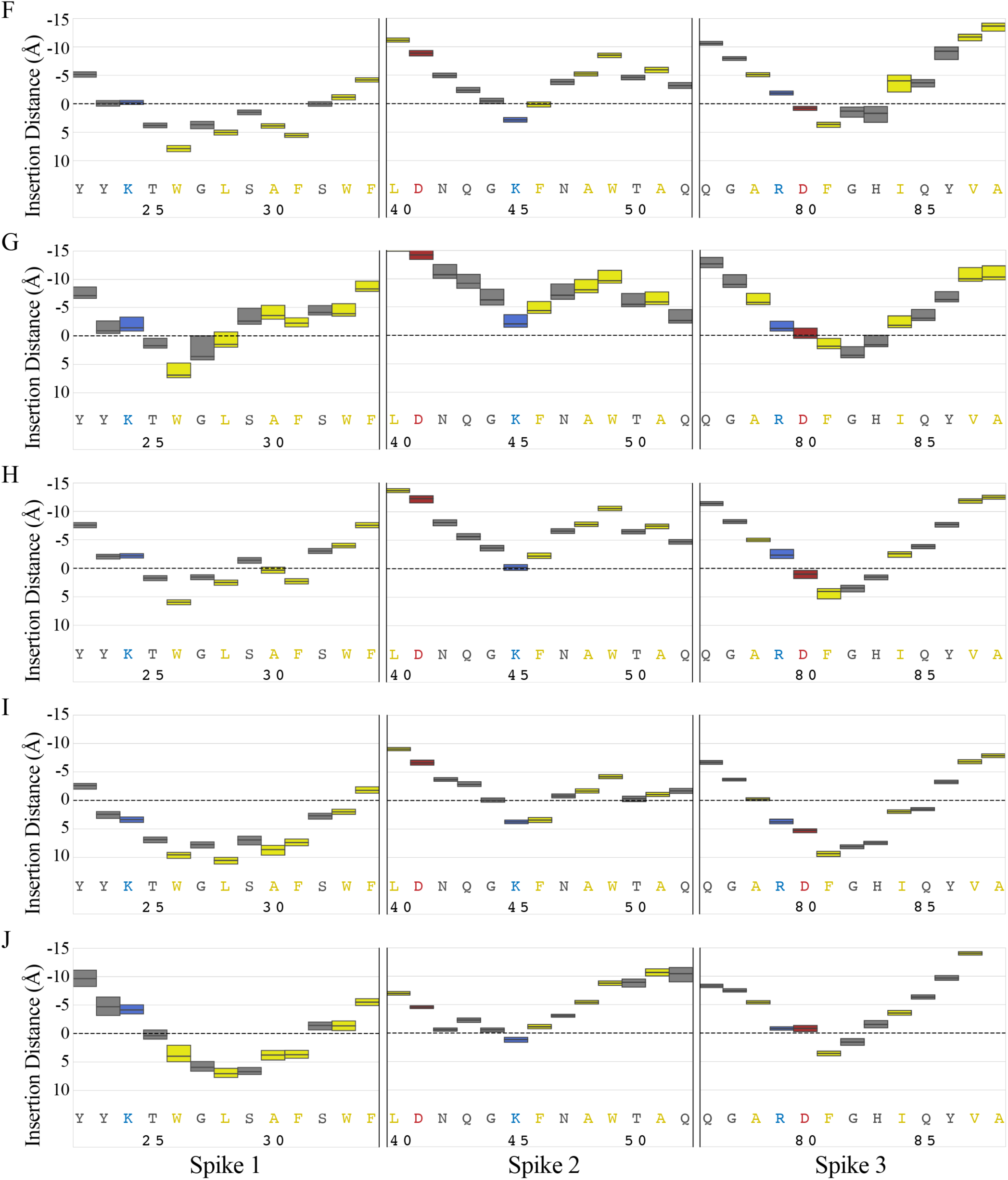
Insertion depth analysis of LactC2 Spikes 1, 2, and 3 in the presence of Ca^2+^. Residue insertion depths are color coded by residue type: polar (gray), non-polar (yellow), basic (blue), and acidic (red). (A-J) Replicas 1-10, respectively.

**Figure S5.**
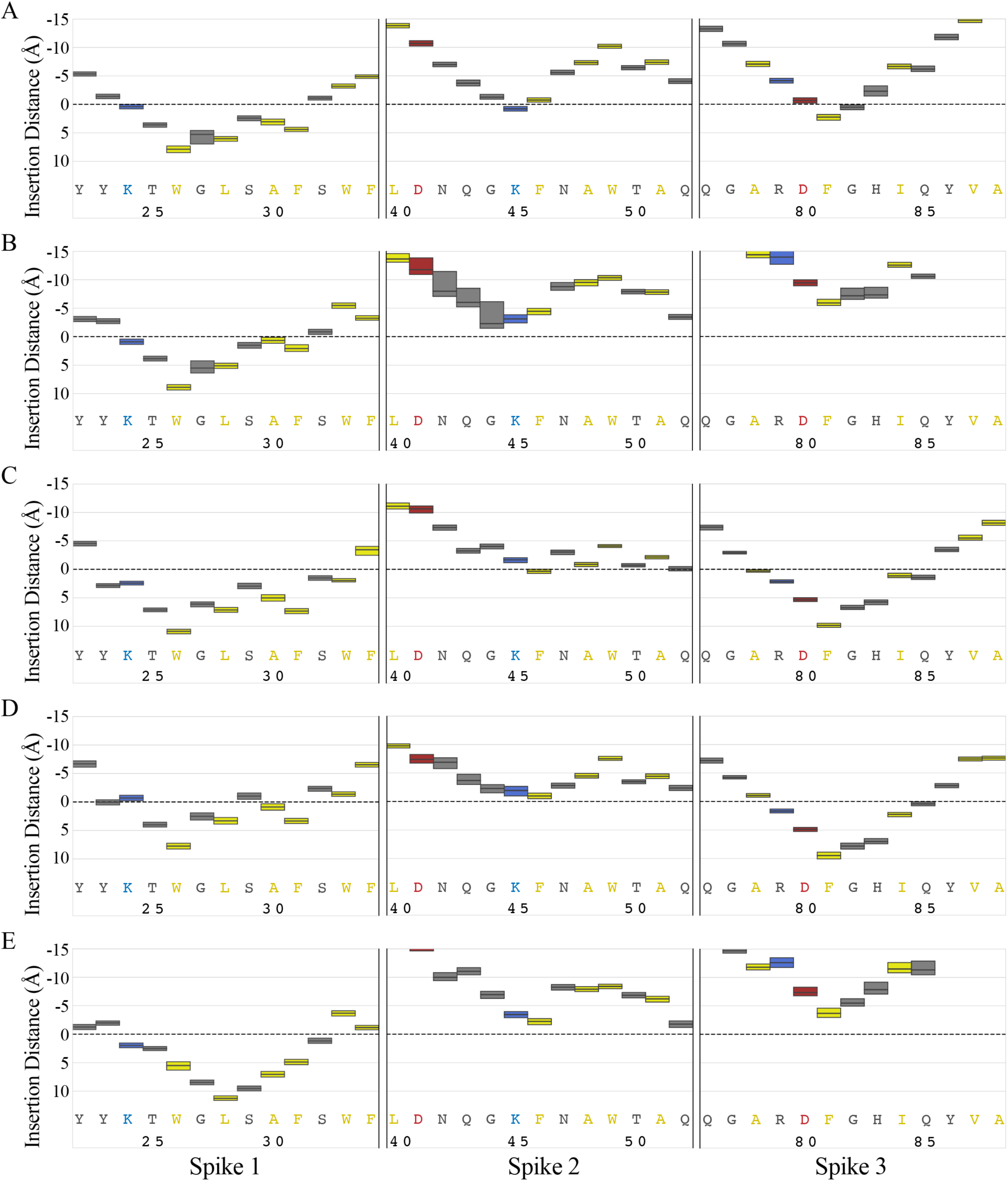
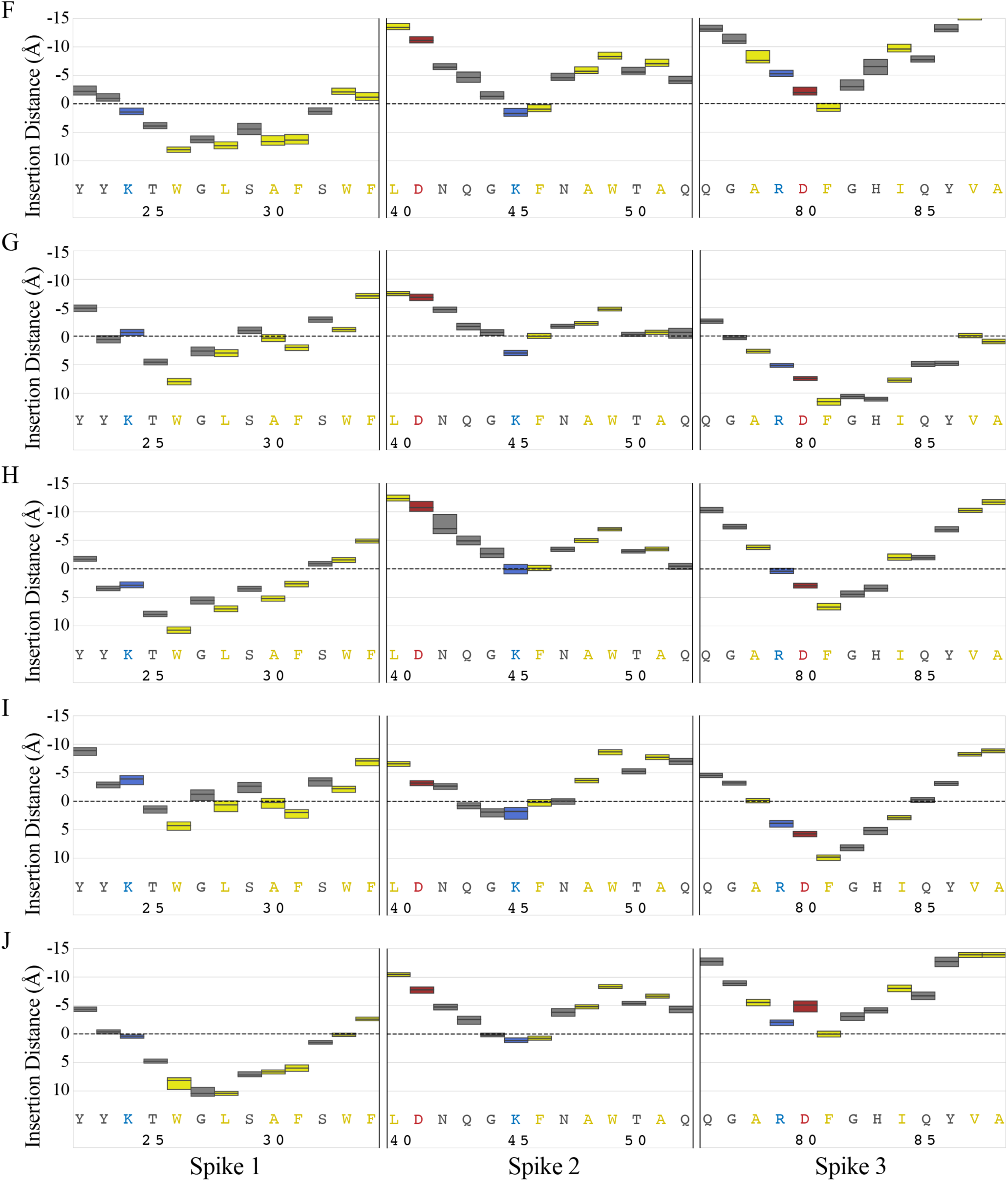
Insertion depth analysis of LactC2 Spikes 1, 2, and 3 in the absence of Ca^2+^. Residue insertion depths are color coded by residue type: polar (gray), non-polar (yellow), basic (blue), and acidic (red). (A-J) Replicas 1-10, respectively.

**Figure S6.**
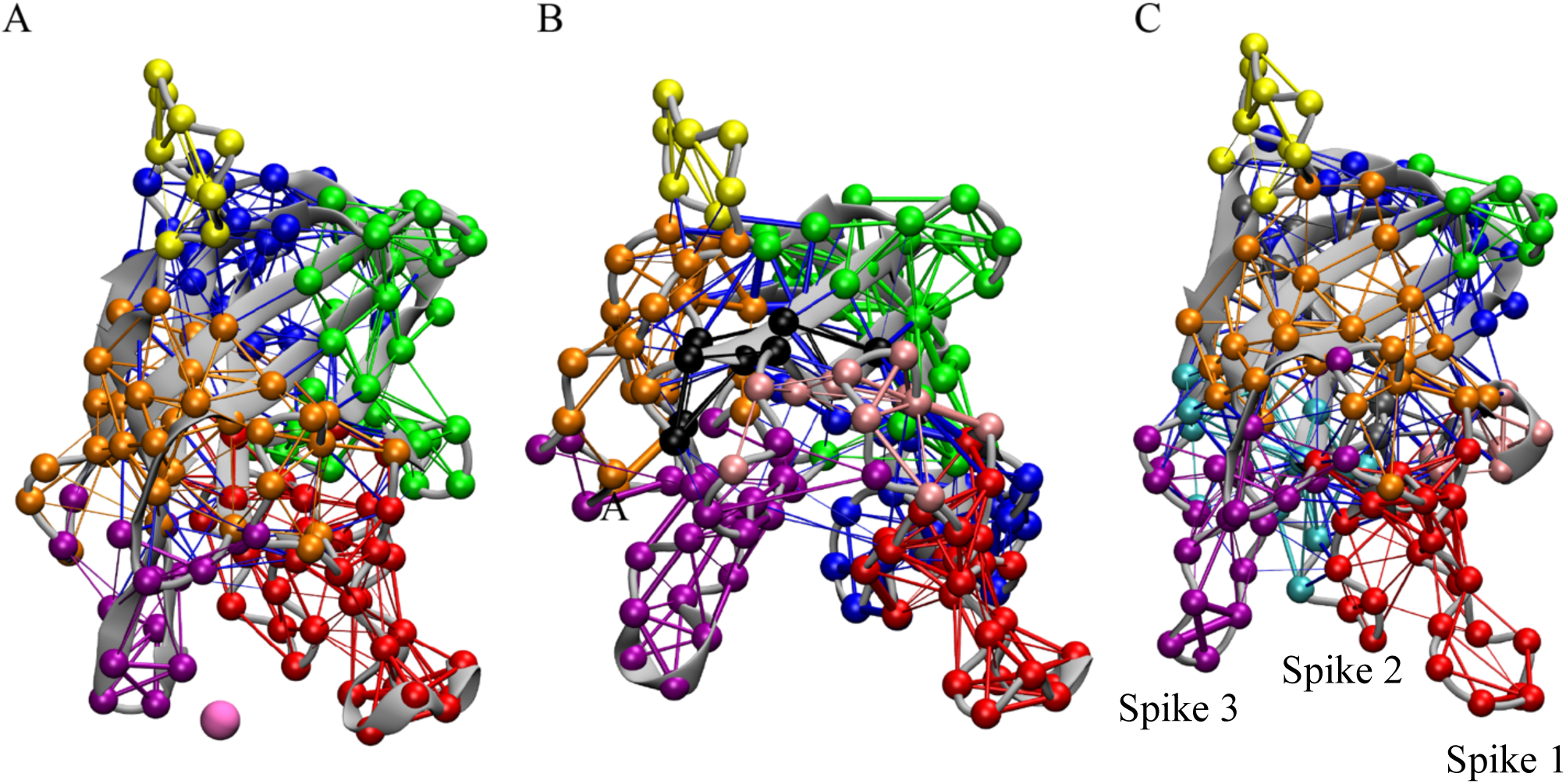
Network analysis of inserted LactC2 domain in (A) the presence of Ca^2+^ (pink sphere), (B) absence of Ca^2+^, and (C) in a water box. The secondary structure of LactC2 is shown in cartoon representation. Network nodes and connecting edges are colored by community (red, blue, orange, yellow, gray, green, purple, light pink, cyan, black).

**Figure S7.**
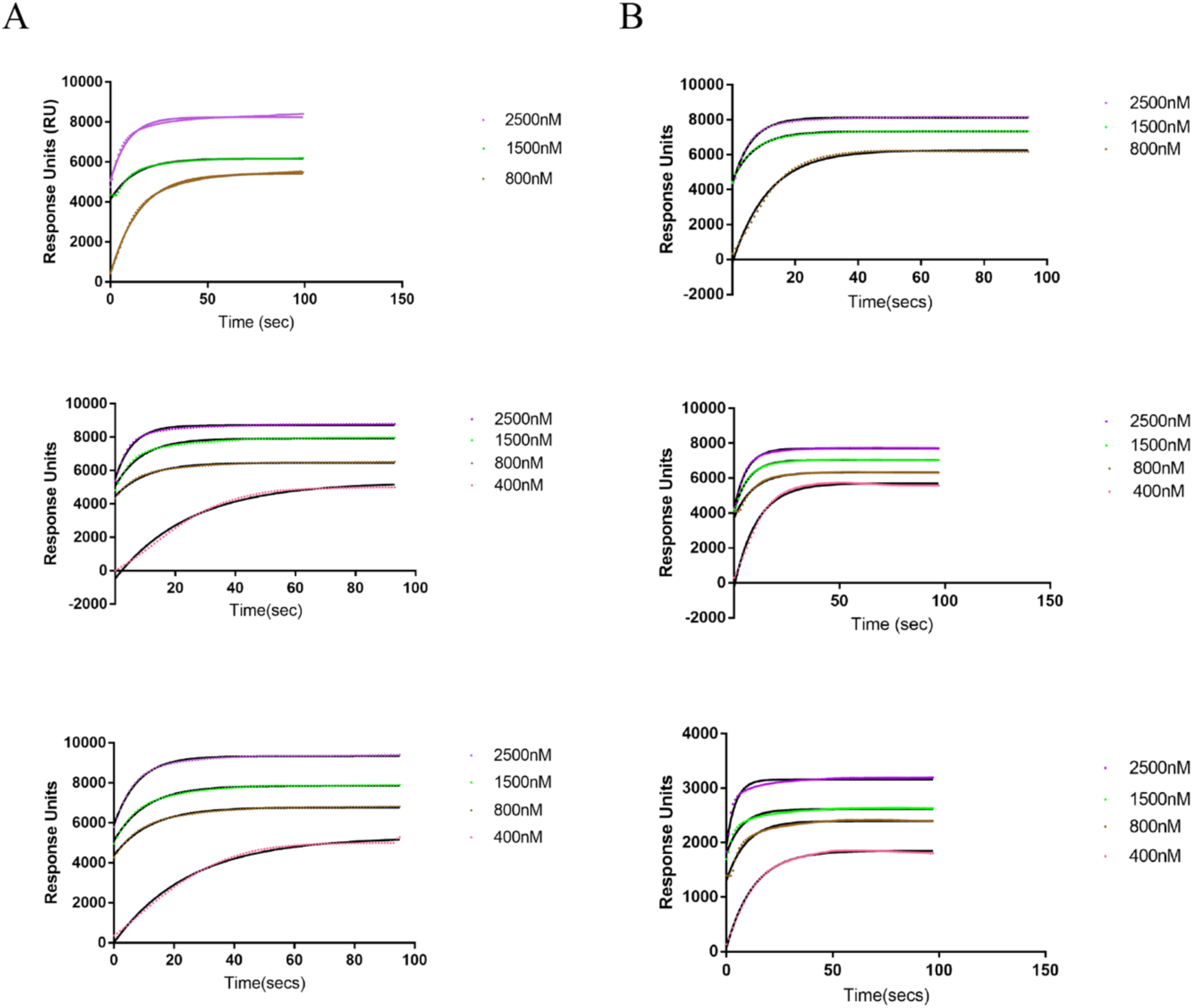
Association studies of lactadherin to nanodiscs both with (A) and without (B) Ca^2+^. Each experiment was completed in triplicate the results from each individual experiment are shown.

**Figure S8.**
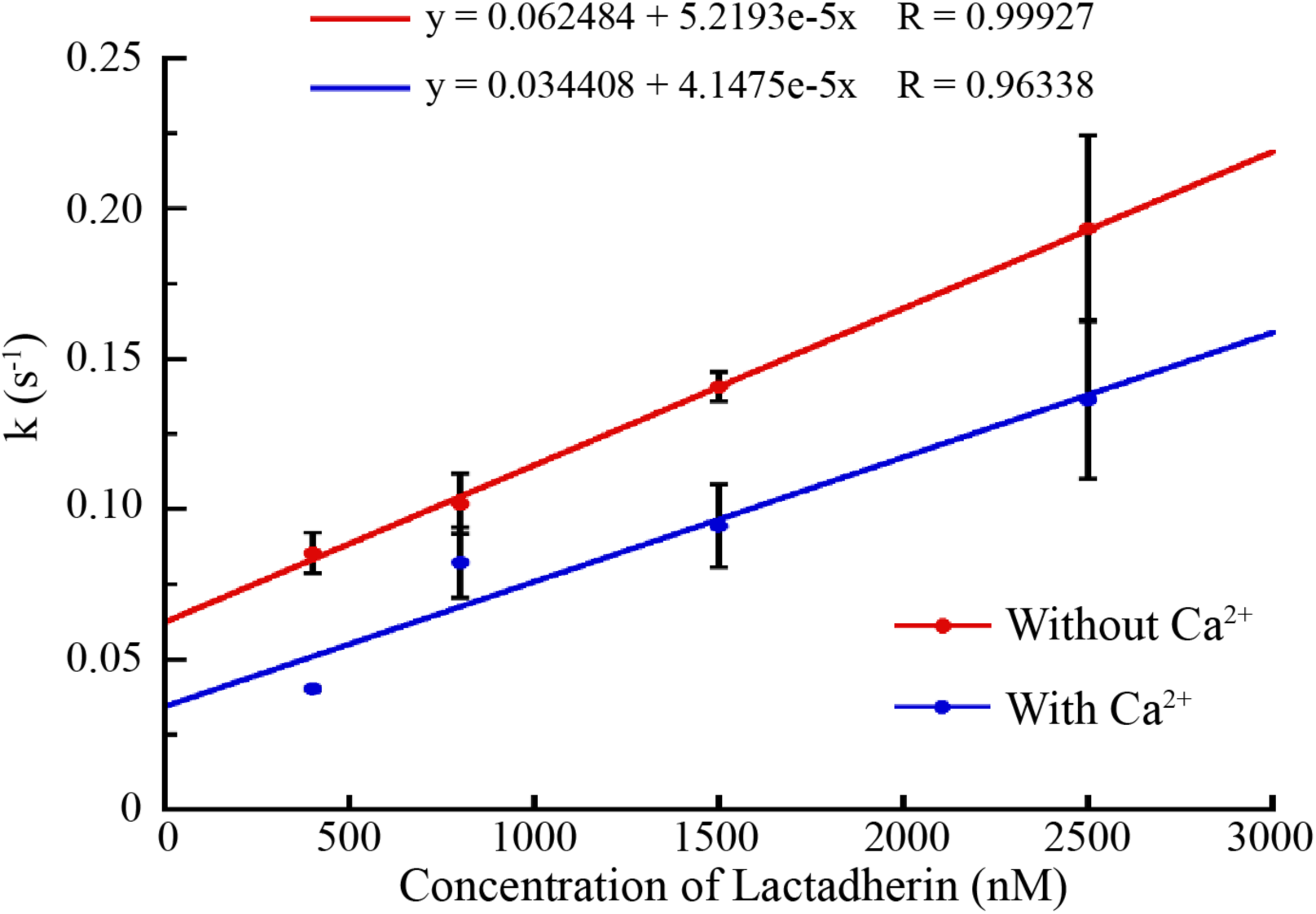
Comparison of association rates (K_on_) of lactadherin to the nanodiscs both in the absence of Ca^2+^ (red) and in the presence of Ca^2+^ (blue). The slope of rate versus concentration lines approximate the K_on_ values.

## Notes

### Competing Interest Statement

The authors have declared no competing interest.

### Summary of Updates

Format was converted into an article. Figures 2-4 revised and SI added.

